# Population genomics perspectives on convergent adaptation

**DOI:** 10.1101/503284

**Authors:** Kristin M. Lee, Coop

## Abstract

Convergent adaptation is the independent evolution of similar traits conferring a fitness advantage in two or more lineages. Cases of convergent adaptation inform our ideas about the ecological and molecular basis of adaptation. In judging the degree to which putative cases of convergent adaptation provide independent replication of the process of adaptation, it is necessary to establish the degree to which the evolutionary change is unexpected under null models and to show that selection has repeatedly, independently driven these changes. Here, we discuss the issues that arise from these questions particularly for closely-related populations, where gene flow and standing variation add additional layers of complexity. We outline a conceptual framework to guide intuition as to the extent to which evolutionary change represents the independent gain of information due to selection and show that this is a measure of how surprised we should be by convergence. Additionally, we summarize the ways population and quantitative genetics and genomics may help us address questions related to convergent adaptation, as well as open new questions and avenues of research.

---

Convergence is the independent evolution of similar features in two or more lineages (Losos, 2011). Examples of convergence have played a fundamental role in our understanding of adaptive evolution (see Losos, 2011, for review). Classically, convergent evolution has been used as one line of evidence for the adaptive value of a trait. When convergence is the result of natural selection producing the same solution in response to repeatedly encountered biotic and abiotic pressures, we treat these as natural replicates of a similar process to understand the ecological and phenotypic bases of convergence (Harvey et al., 1991) and the molecular basis of evolution (Stern, 2013; Martin and Orgogozo, 2013; Rosenblum et al., 2014; Storz, 2016).

The broad definition of convergence permits us to think of convergence occurring at multiple levels. At the broadest scale, convergence in function may occur in species facing similar ecological challenges, consistent with view that there are sometimes many phenotypic ways to achieve a similar function (Wainwright, 2007). For example, carnivorous plants have evolved similar strategies to adapt to their low-nitrogen environments but have done so via many distinct morphological and physiological mechanisms across five orders of flowering plants (Givnish, 2015). Convergence may also occur on the level of a specific phenotype. Carnivorous pitcher plants have striking convergence for trap morphology (Thorogood et al., 2018). When convergent phenotypes arise in highly-diverged species, it has often been presumed that these changes are due to divergent genetic mechanisms (see Stern, 2013, for review). However, perhaps surprisingly, there is increasing evidence that some phenotypes often converge due to selection on similar molecular mechanisms, with selection occurring at the same gene or even the same mutational change (Christin et al., 2010; Martin and Orgogozo, 2013; Stern, 2013). Again within carnivorous plants, Fukushima et al. (2017) find convergent amino acid substitutions in the evolution of digestive enzymes across four species. Despite these distinctions, all levels can be encapsulated in the term convergence. Additionally, following Arendt and Reznick (2008), the single term can be used to describe both parallelism and convergence, regardless of the phylogenetic scale, ancestral state, or whether the underlying genetic mechanisms are the same or different. In this perspective piece, we discuss how adaptive convergence at the level of phenotypes, genes, and allele frequency change can be studied with population genomic data. In establishing a case of convergent adaptation two main conditions must be satisfied. First, the character must be adaptive. That is, it enhances the fitness of the organisms that bear it, relative the ancestral state (Futuyma, 2013), with many definitions requiring adaptations to be the product of natural selection for their current role (Gould and Vrba, 1982; Baum and Larson, 1991). Second, the adaptive changes in the different populations must be independent from each other. Below, we expand upon these conditions and show how they might be quantified through measures of the independent work done by natural selection in different populations. Throughout this article, we focus on how population-level genomic data both forces us to grapple with the issues that arise in identifying convergent adaptation among closely-related taxa and provides novel ways to approach these and related questions.

## Identifying adaptations

Many different lines of evidence from developmental biology to field studies must often be brought together to demonstrate that a character is an adaptation. Often, one of the first steps in assessing whether some set of convergent characters are adaptations is to establish that the changes observed are unexpected by chance (i.e. under simple null evolution models). To address this problem for convergence at macro-evolutionary scales, phylogenetic null models and methods have been developed both for discrete and continuous traits (Zhang and Kumar, 1997; Stayton, 2008; Castoe et al., 2009; Mahler et al., 2013; Thomas and Hahn, 2015; Yeaman et al., 2018), as well as for testing the association of a character with some specific ecological or environmental variable (Harvey et al., 1991; Harmon, 2018). Convergence changes that are not due to the vagaries of divergence or genetic drift are good candidates for adaptations. However, even if the convergence is unexpected, this does not necessarily mean that it is adaptive. For example, it could be due to developmental constraint or releases from constraint in similar ecological environments. Therefore, it is important to provide evidence that the changes are associated with increased fitness in the relevant environments. One way of doing this is to provide evidence that directional selection repeatedly drove the convergent change. The field of population genetics has a long history of discerning phenotypic and allele frequency shifts due to selection from those due to neutral processes. In this article, we review some of the ways these approaches can be used and extended to studying cases of convergence.

## Establishing independence

Showing that changes observed were truly independent is necessary to gain insight about adaptation from studies of convergence; otherwise, they may just represent a single replicate of the adaptive process. Establishing that changes were truly independent is difficult. Even with a well-resolved phylogeny, with no gene flow and nodes with sufficient temporal separation such that there is no incomplete lineage sorting (ILS), secondary loss must be distinguished between convergent evolution to show that a trait is not shared from a common ancestor (Wake et al., 2011). Establishing independence for more closely related taxa can be even more challenging. Closely-related populations may be more likely to share independently derived states that were absent or rare in their ancestor. This may occur not only through independent mutations recurrently becoming a substitution in different parts of the phylogeny, but from shared variation present in their ancestor and introgression. These latter modes are expected to be more prevalent between closely-related taxa because of their higher likelihood of interbreeding and retention of ancestral polymorphism.

The sharing of traits or alleles incongruent with the population or species tree due to ancestral variation (i.e. ILS shown in Figure 1A) has been termed hemiplasy (to contrast it with homoplasy 2008; Wake et al., 2011). This term has also been more generally applied to alleles shared by gene flow and with some authors seeing it as an alternative explanation to convergence (Hahn and Nakhleh, 2016; Mendes et al., 2018; Guerrero and Hahn, 2018). While these approaches to assess discordance between gene trees and species trees that lead to spurious inference of convergence are important, we argue that the sharing of an allele due to ILS or introgression does not invalidate the case for convergent adaptation. Rather, these questions are focused on identifying the level of independence. When convergence at the level of the same molecular substitution between distant species is observed, we can usually assume that this was due to multiple, independent mutations (i.e. convergent mutations). However, the key question is whether selection has independently increased the allele to fixation multiple times (i.e. there has been convergent substitutions). Taxa sharing the same substitution due to ILS or gene flow means there has not been independence at the level of the mutational change; however, the allele frequency change at the locus can still be independent, that is convergent, across populations. For example, following populations 1 and 3 in Figure 1A, even if an allele was shared due to ILS, selection may have repeatedly driven up the allele in each population separately.

**Figure 1:**
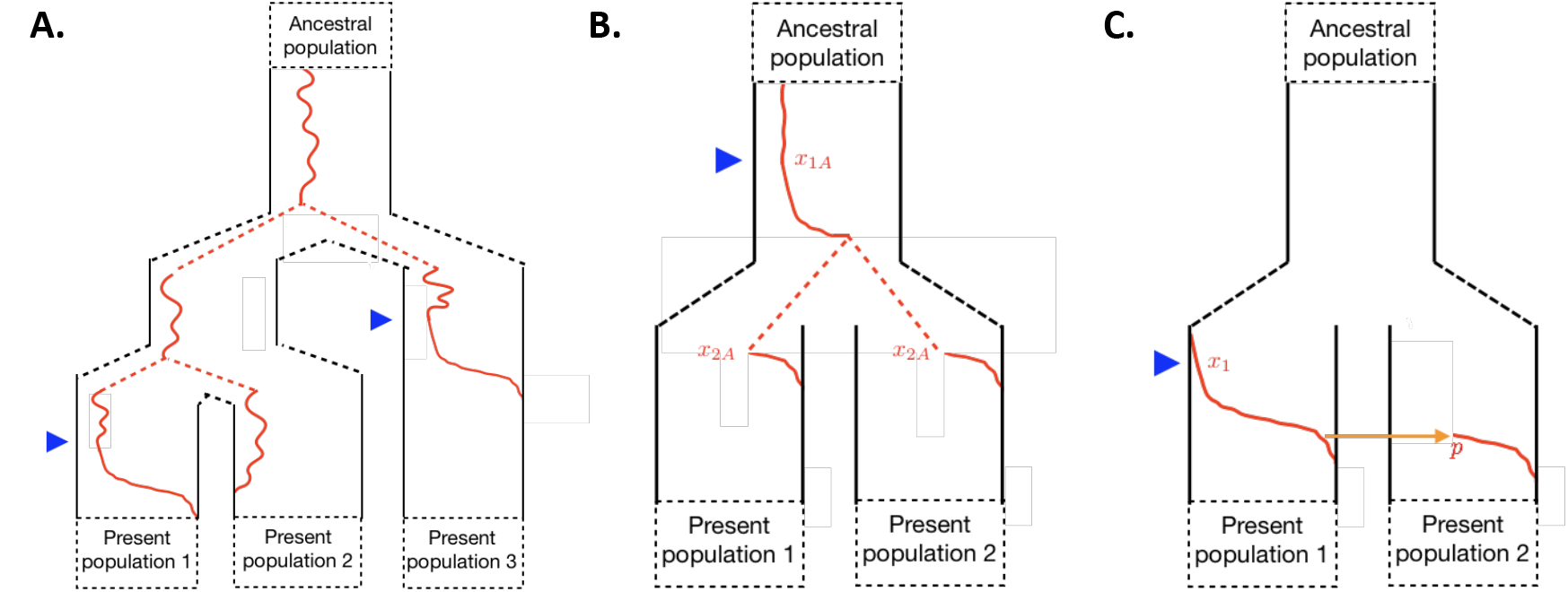
Allele frequency trajectories (red) at a single locus for two or more populations. A) The allele was standing in the ancestor of three populations where it was fixed in populations 1 and 3 but lost in population 2. This case of incomplete lineage sorting (ILS) may falsely resemble independent mutations in populations and 3, but may still be convergent adaptation if selection for the allele occurred twice independently (e.g. onsets of selection marked by blue triangles). B) The beneficial allele was standing in the ancestral population of our present-day populations. Selection started when the ancestral allele frequency was *x_1A_* and increased in frequency such that it was at *x_2A_* when the populations split. Then, it continued to increase in both daughter populations to fixation. C) The beneficial allele arose in a single present-day population and spread via introgression into the other population. The allele was at frequency *x*_1_ at the onset of selection and migrated into the recipient population at frequency *p* where it also increased in frequency to fixation.

In highlighting levels of independence, it is not our intention to add more vocabulary to a perhaps already complex area. Instead, we aim to make clear the questions we seek to address when studying convergence at various levels. We, in particular, are interested in dissecting these levels of independence in cases of convergence at the same gene (Lee and Coop, 2017), to gain insights about adaptation into how a particular variant or haplotype offers a selective benefit. For example, seeing different mutations is informative about the mutational target for adaptation. Yet, the repeated use of standing variation, especially if those alleles are very rare, still informs us about the mutational target size. An example of both instances exist in marine populations of threespine stickleback that have repeatedly, independently have adapted to freshwater environments via changes at the EDA and PITX1 genes. The fact that many independent mutations at PITX1 underlie adaptation (Chan et al., 2010; Shapiro et al., 2004) suggests that this state is relatively accessible to the population by mutation or that other possible mutations may be too constrained by pleiotropy. Conversely, the EDA haplotype, which had been standing and reused many times by adaptation in freshwater sticklebacks (Colosimo et al., 2005), is suggestive that populations cannot access a similar, relatively low pleiotropy state more rapidly by mutation. The frequency of these different modes of convergence speaks to the extent to which populations are mutation-limited and to the role of standing variation and gene flow in adaptation (Welch and Jiggins, 2014). Thus, regardless of the precise terminology we use, it is important to keep clear statements about the level of independence and the role selection has played in these changes. In this article, we introduce a conceptual framework to quantify the degree to which allele frequency shifts have been independent. We then review current approaches in population and quantitative genetics and genomics and how those can be extended to include more explicit tests for independence.

## Quantifying the independence of convergent adaptation, the accumulation of information, or when to be surprised

One conceptual way forward in quantifying convergent adaptation may be to assess the degree to which selection has independently driven phenotypic and allele frequency change. Quantifying the work done by selection among taxa can help to guide intuition as to when to be impressed by convergence. One reason why adaptive convergence among lineages that are phylogenetically distant is surprising is that these are clear examples of selection independently, repeatedly reaching similar states that would be unlikely under chance or drift alone. One measure of this is the extent to which selective deaths (or births) underlying an adaptation in two or more populations are independent. For example, if light colored pelts are an adaptation to artic environments, we have high confidence in calling the light colored pelts of arctic foxes and hares as convergent adaptations since the individuals who died due to predation pressure, that drove the adaptation via allele frequency change, were clearly different sets of individuals (being foxes and hares, respectively).

More formally, we can calculate the number of selective deaths needed to generate a given allele frequency change. In a population of size *N*, the selective deaths imposed under a deterministic, directional selection model where the fittest homozygote has selection coefficient *s*, move an additive allele from frequency *x_1_* to *x_2_* in *t*_1_ to generations is *NL* where *L* is

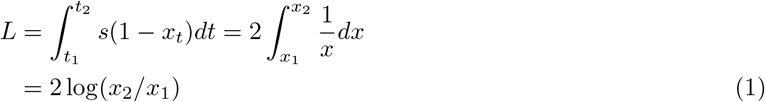

as originally derived by Haldane (1957) in his discussion of the substitution load. (We return to the issues with Haldane’s load argument below.) We will focus on fixed alleles such that

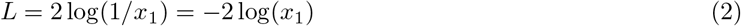

This measure of the work done in fixing an allele due to selection does not depend on the strength of selection since it takes a longer time to fix a more weakly selected allele than a strongly selected allele so the same total number of selective deaths are incurred. Additionally, most of the work done by selection to increase fitness occurs when the allele is very rare in the population. When the allele becomes common in the population, the majority of individuals carry the allele so fewer individual deaths are due to differences in the genotype at this locus. Under this measure of work done by selection, we should be much more impressed by selection that repeatedly moving an allele from 10^−4^ to 50% than 50% to 100%, despite being comparable frequency changes and taking roughly the same time under an additive model.

We can extend this idea to obtain a metric of the proportion of the work done by selection that is shared between populations that both have experienced allele frequency change at the same locus. Here, we apply this to two cases shown in Figures 1B and 1C. We first study the case that a selected allele has increased in frequency in the ancestral population from *x_1A_* to *x_2A_*, and following a population split increased from *x_2A_* to fixation in both of the daughter populations (see Figure 1B). Here, the work done by selection in the ancestral population (*L_A_*, i.e. that shared between our populations), the work independently done in the two daughter populations (*L_D_*), and their ratio can be written as

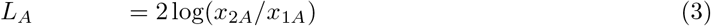

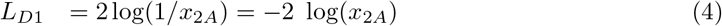

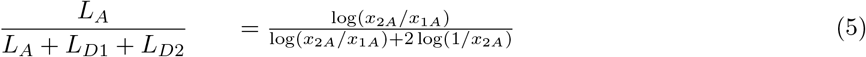

Thus, if the allele selectively increased from a low to intermediate frequency in the ancestral population before the split (*x_1A_* is ⪡ *x_2A_*), much of the work done by selection is shared between the daughter populations. In the absence of other information, it is parsimonious to assume that 100% of the selective deaths are shared between the two sister populations and that the allele fixed in the ancestral population (*x_2A_* = 1). This agrees with our intuition and ideas of phylogenetic parsimony that, in the absence of other evidence, all of the change occurred only once in the ancestral population. That is, we should not infer the repeated work of selection where it is unnecessary. This measure also emphasizes that we should be much more impressed by repeated adaptation from very rare standing variation than common variation. Similarly, in the ILS example shown in Figure 1A, in the absence of other information it is parsimonious to assume that the that allele fixed in populations 1 and 3 was relatively common in the ancestral population; so, its repeated fixation by drift was not unlikely. Only in the case where we have other information, for example that neutral ILS is very rare given the timespan involved, do we have some evidence that convergence is unusual.

As a final scenario, we consider the case when a selective allele increases in frequency in one population (1) due to selection before being introduced into a second population (2) by gene flow and fixing in the second population (see Figure 1C). For simplification, we assume a single pulse of admixture that introduces the allele into the second population at frequency *p*. The work done by selection in populations 1 and 2 are *L*_1_ = –2log(*x*_1_) and *L*_2_ = –2log(*p*), respectively, for a total load of *L_T_* = –2(log(*x*_1_) + log(*p*)). Therefore, the proportion of selective deaths that are shared is

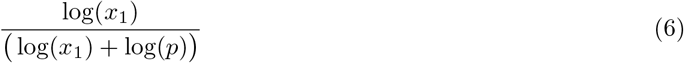

Since most of the selective deaths involved in fixing an allele occurs when the allele is rare, they are shared between populations unless the frequency that the allele is introduced into population 2 is small (i.e. *p* close to *x*_1_). For example, if the allele fixed from an initial frequency of 1/1000 in population 1 and admixture into population 2 initially pushed allele to 10% frequency, a large portion, 75%, of the work done by selection is shared.

We can also relate this measure to the amount of information gained independently in different populations. As noted by Kimura (1961), the load involved in fixing a single substitution (Equation 2) is twice the log of the probability of a neutral allele fixing from *x_1_* and so is a measure of the information accumulated by selection as compared to a neutral model (in an information theoretic sense, Shannon, 1948; Cover and Thomas, 2012). We explain this connection in Appendix A.1 and how it follows from the fact that Equation 2 can be seen as the average log-likelihood ratio of the probability of an allele fixing under the selective model compared to the neutral model. This information gain under a model of selection compared to neutrality is a measure of the unlikeliness of the state of the system encoded by selection as compared to a neutral model. Extending this argument in Appendix A.1, we show that the decomposition of the shared and independent work done by selection through selective deaths, e.g. Equations 3–6, corresponds to the shared and independent gain of information due to selection. Specifically, we can express the additional work done by selection in a second population in terms of how improbable the allele frequency change was in a second population under a model of genetic drift, given the change already achieved in the first or ancestral population. We also show how this measure of convergence, as the independent work performed by selection, can also be expressed as the statistical ‘surprise’, under a neutral model, as observing what has been achieved in a second population conditional on what happened in a first or ancestral population. While we have considered just two populations here, we can extend this framework to larger numbers of populations. However, the main insight, that we want to see repeated large allele frequency changes from initially rare alleles, is clear from just the two population case.

This framework may be useful in aiding our intuition about the degrees to which selective events have been independent and so can be considered to be convergent. At the macro-evolutionary scale, with no gene flow or ILS, convergent substitutions in different taxa can be thought of as completely independent allele frequency changes. Thus, seeing a specific, selection-driven substitution twice offers double the information gain by selection. It seems natural to then think of simply summing up the information gained over multiple substitutions across loci. However, one issue with this approach is that we cannot simply sum the work done by selection (i.e. selective deaths) across multiple loci, unless the loci contribute multiplicatively to fitness. The same information content can be accumulated for far fewer deaths under negative fitness epistasis (e.g. most efficiently due to truncation selection). This argument forms the basis of the rejection of Haldane’s substitution load argument, and Kimura’s gain of information extension, as a limit on the rate of evolution (Maynard Smith, 1968; Sved, 1968; Felsenstein, 1971). Combining information over loci additively, that is assuming multiplicative epistasis, seems a fine first approximation, but this limitation should be borne in mind. However, pushing beyond that limitation the information gained through selection is a more general concept of a statistical measure of the work done by selection in pushing evolution towards highly improbable states. A number of authors have explored the idea of statistical information as a measure of work done by selection and more broadly the role of selection in decreasing the entropy of states including in the presence of different forms of epistasis (Iwasa, 1988; Watkins, 2002; Peck and Waxman, 2010; Mustonen and Lassig, 2010; Barton, 2017). These ideas extend naturally to quantitative traits (Iwasa, 1988; Frank, 2012; Barton, as well as substitution models from molecular evolution (Iwasa, 1988). The null models that we judge our gain of information due to selection against can also include mutation and mutational biases. Therefore, convergent adaptation, at the genetic or phenotypic level, can potentially be conceptualized in this way as the independent gain of information due to selection.

## Using population genetics to identify convergent adaptation

Population and quantitative genetics and genomics can provide evidence for convergent adaptation both by helping to establish the degree to which evolutionary changes among populations are independent and by providing evidence that, historically, individuals with a given allele had enhanced fitness relative to the ancestral state. In this section, we review how single locus and quantitative trait tests, and haplotypic or linked variation data, can be used to test the conditions of convergence.

### Single locus and quantitative trait tests

In evolutionary studies of characters across species, comparative methods (Felsenstein, 1985; Harvey et al., 1991) make use of the phylogeny to judge whether the sharing of a character among taxa is expected under some null model (e.g. independent evolution down a phylogeny) and whether characters are correlated with some ecological variable beyond null expectations (see Harmon, 2018, for a recent review). For example, in phylogenetic mixed models, the phenotypes of a set of species is regressed on some environmental variable as a fixed effect and divergence along the phylogeny is incorporated as a random effect, with the covariance structure of the random effect being specified by the matrix of shared branch lengths (Lynch, 1991; Housworth et al., 2004). Comparative methods provide a series of tools for evaluating convergence for discrete and continuous characters and use convergence to gain independent information to test evolutionary hypotheses.

There is an equivalent body of work in population genetics which involves shorter evolutionary time scales. In answering these questions on short-time scales, one equivalent to the phylogeny is the relatedness or kinship matrix (Hadfield and Nakagawa, 2010; Stone et al., 2011). There are different ways of defining this matrix, but broadly the elements of this matrix specify the amount of sharing of alleles between two individuals or populations, above that expected in a reference population (often some hypothetical ancestral population or the total sample), due to shared coancestry and genetic drift (Goudet et al., 2018). This is conceptually similar to how a species phylogeny specifies the shared branch lengths for pairs of species back to their common ancestors and hence the degree of shared ancestry relative to the most recent common ancestor of all the samples. However, using the relatedness matrix is a somewhat different statement from the importance of species phylogeny in judging trait similarity since introgression and ILS increase the chance that alleles and traits can be discordant with a species phylogeny (Hahn and Nakhleh, 2016; Mendes et al., 2018; Guerrero and Hahn, 2018). This issue is well accommodated for by using the relatedness matrix which naturally accounts for allele sharing that is discordant with the species tree. An additional benefit is that models based on the relatedness matrix can be used in situations where population relationships are not well described by a simple branching phylogeny, for example isolation by distance, which is frequently encountered when dealing with populations separated by short timespans.

Tests of selection involving an empirical relatedness matrix have become more common with the increase in data from large numbers of closely-related populations. For example, the association between allele frequencies at a locus and an ecological variable can be assessed by using the relatedness matrix to model the null expectations of populations that vary and covary in their deviations due to genetic drift (Hancock et al., 2008; Coop et al., 2010; Frichot et al., 2013). A similar body of work tests for non-neutral population differentiation while accounting for population structure by using the relatedness matrix or the eigen-vectors (principal components) of this matrix Bonhomme et al. (2010); Günther and Coop (2013); Duforet-Frebourg et al. (2015); Galinsky et al. (2016). By incorporating the relatedness matrix as a null model into population-genetic tests, shared population history and gene flow is naturally accommodated for in assessing environmental correlations or non-neutral allele frequency divergence. While these tests use the independent evidence of allele frequencies across populations to test for deviations away from a null model, they do not generally test explicitly for adaptive convergence. This is because they are not usually used to take the extra step of determining whether similar patterns of non-neutral allele change in distinct subsets of the species range are in fact independent or can be explained by selection-driven allele frequency change in one region followed by gene flow and drift spreading the change to other regions. However, researchers do already informally test for convergence of allele frequency change across species ranges, for instance in studying parallel selective clines on multiple continents or at smaller geographic scales. (See, for example, the work on *Drosophila* genomic clines of Adrion et al. (2015); Sedghifar et al. (2016); Machado et al. (2016).) One possibility to formalize a test of convergence is to condition on the observed allele frequency change in one subset of the species range, using the relatedness matrix, while testing for non-neutral allele frequency change elsewhere. Additionally, frameworks that use admixture graphs, parameterized versions of the relatedness matrix (Pickrell and Pritchard, 2012), to determine whether multiple instances of selection on distinct branches are being developed to examine allele frequency change across populations (Refoyo-Martínez et al.,. Therefore, we are now in position to move toward formally testing for non-neutral convergence in allele frequency changes among populations.

In studying phenotypes, we can also address similar questions across closely-related populations. Quantitative geneticists have long asked whether common-garden phenotypes are too variable across populations compared to the allele frequency differentiation (often using *Q_ST_-F_ST_* comparisons; see Whitlock, 2008; Leinonen et al., 2013, for reviews). These tests were originally restricted to relatively simple models of population structure. Recently, there has been a move to using kinship matrices to test for non-neutral divergence in phenotypes or polygenic scores across populations, allowing shared neutral trait divergence due to shared population history and gene flow to be more fully accounted for (Ovaskainen et al., 2011; Karhunen et al., 2013; Berg and Coop, 2014; Berg et al., 2017; Josephs et al., 2018). These tests are predicated on the idea that the additive genetic phenotypic variation among populations is the sum of allele frequency changes at many loci. Therefore, its neutral properties are well captured by the relatedness matrix in a conceptually similarly way to how Brownian-motion models capture divergence due to phenotypic effects of many small substitutions on a phylogeny. These relatedness-based models can be used to ask whether there is support for phenotypes covarying with environments more than expected under models of genetic drift (Karhunen et al., 2013; Berg and Coop, 2014; Berg et al., 2017) and to assess non-neutral divergence along different axes of population structure. These approaches have been used to quantify the degree to which non-neutral genetic phenotypic change among populations is independent versus a consequence of shared history. For example, assessing how unexpected the phenotypes or polygenic scores of one set of populations are, conditional on another distinct subset (Berg and Coop, 2014; Berg et al., 2017). These ideas have also been extended into explicit tests of non-neutral phenotypic or polygenic score change on admixture graphs (Racimo et al., 2018) that could be used to detect repeated changes among populations.

In conclusion, methods to assess non-neutral, repeated evolutionary change across many populations are rapidly being developed. These developments mirror many of aspects of phylogenetic comparative methods, but extend them to situations where population history is not well represented by a single tree. In moving forward, we need to more explicitly test for convergence and incorporate more intuitive measures of when this convergence is surprising.

### Linked variation

Beyond single locus tests or tests at the level of the phenotype, knowledge of the population genetic variation at set of linked sites helps address questions related to convergence that may not be possible otherwise. As discussed above, convergent evolution at the level of a gene or allele is most surprising when selection repeatedly and independently pulls alleles up from rare frequency. By comparing haplotype patterns across populations, we can hope to build evidence towards demonstrating this as well as establishing independence to identify convergence.

One common signal leveraged in population genomic scans for selection is the hitchhiking effect. The rapid increase of a rare, beneficial allele gives little time for mutation and recombination to introduce genetic variation on the haplotypic background of the allele, resulting in a decrease in allelic diversity in this region (Maynard Smith and Haigh, 1974; Kaplan et al., 1989). Thus, a strong signal of a selective sweep at a locus is evidence that selection has led to a gain information in the population. With have multiple populations exposed to some novel selection pressure, linked variation and haplotype patterns not just within but among populations can be informative. Strong allele frequency differentiation among populations can be used as additional evidence of selection which can be linked to the specific selection pressure driving the change. However, seeing the signal of a selective sweep across several populations does not ensure that selection was independent. For example, there may have been recent selection for the trait once in the ancestor of the present-day populations (i.e. the scenario in Figure 1B where the sweep went to fixation fully in the ancestor or the allele frequency change from *x_2A_* to fixation was due to drift). Additionally, there may have been selection in a single population and sufficient gene flow and genetic drift to increase the frequency of the selected haplotype with no additional selection in the recipient populations (i.e. Figure 1C where now the increase in the recipient population was not due to selection in population 2 but only migration pressure and drift).

Population genomic data helps dissect whether sweeps across multiple populations are indeed true examples of convergent adaptation. If, for example, populations adapt from different or independent mutations within the same gene or different genes that arose and swept, little overlap is expected in the haplotypes among these populations (Figure 2). This builds support for the case that adaptation was likely independent and that information gain due to selection was compounded across populations. Convergent adaptation due selection on the same mutation, i.e. independent adaptive allele frequency change on the same variant (independent substitutions but not independent mutations), may be also distinguishable from cases where most of the change is due to shared selection. In Lee and Coop (2017), we developed an approach to distinguish between the ways convergent adaptation can arise using neutral allele frequency data from sites linked to beneficial loci. Specifically, we derived coancestry coefficients (terms proportional to the relatedness and kinship) both within and between populations experiencing the same selective pressure based on coalescent probabilities. We then used these terms to calculate the composite likelihoods of observing a set of neutral allele frequencies given a specific scenario of selection and selected allele sharing. Previously, we aimed to distinguish among convergent adaptation due to convergent mutations, independent selection on shared ancestral variation, and independent selection on an introgressed variant, as well as a neutral model with no selection. In Appendix A.2, we derive similar coancestry coefficients for the scenario where there has been a complete sweep in the ancestral population to exemplify how this approach can be extended to model scenarios that are not truly convergent adaptation in hopes to distinguish between these signals as well. Other authors have explored models and developed methods for sweeps from standing variation (Peter et al., 2012; Roesti et al., 2014; Berg and Coop, 2015), sweeps from selected alleles spread by migration or introgression (Slatkin and Wiehe, 1998; Santiago and Caballero, 2005; Bierne, 2010; Kim and Maruki, 2011), and ancestral sweeps (Racimo, 2016). We hope that our approach, which combines insights from many of these models into a single framework, can aid investigators in identifying, and distinguishing among models of, convergent adaptation. Additionally, since our approach does not assume an underlying population tree, this may be particularly useful for closely-related populations that may have short branching times and recent or ongoing gene flow.

**Figure 2:**
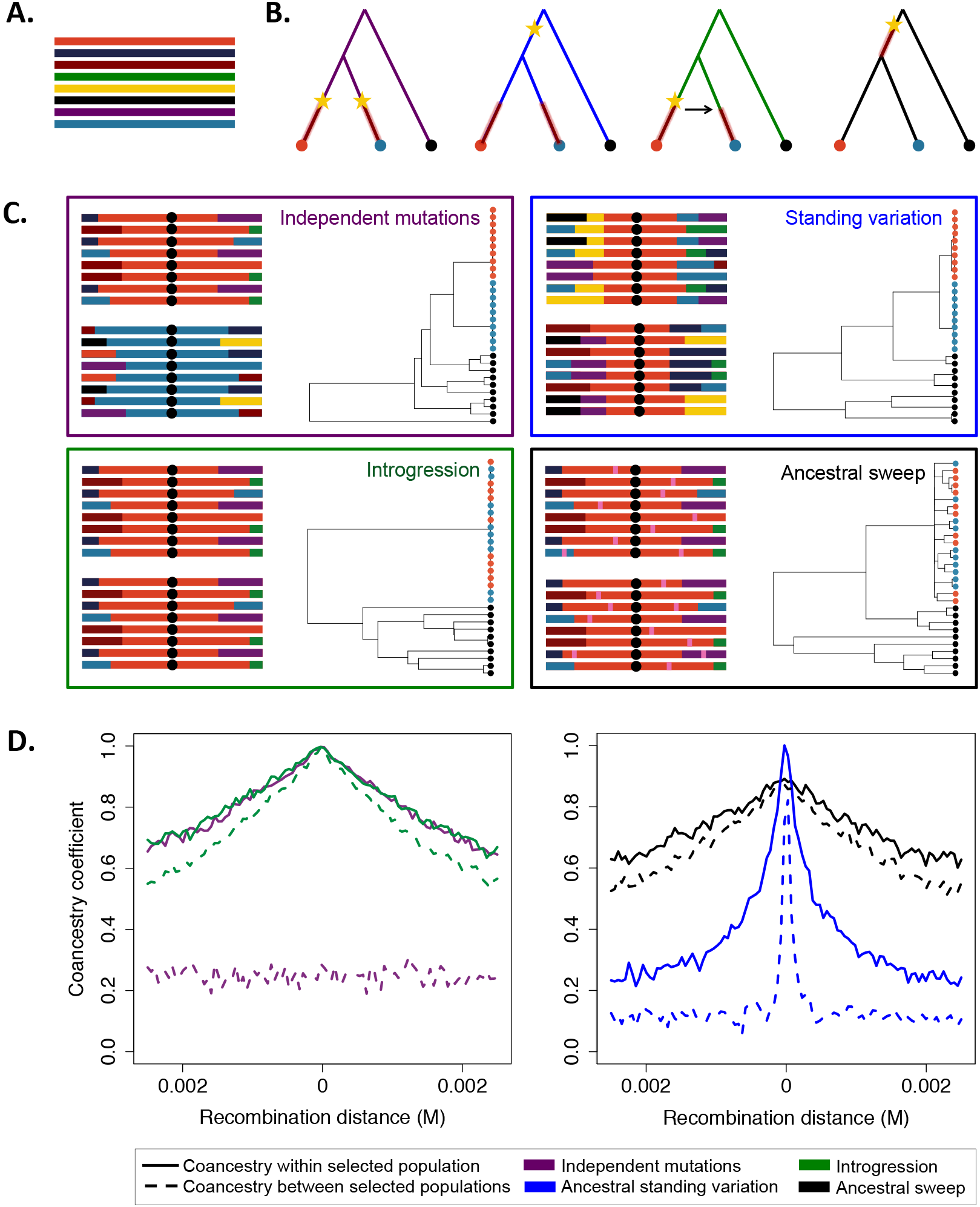
Patterns generated under scenarios of convergent adaptation and not truly convergent (i.e. not independent) adaptation. A) Haplotypic diversity in the ancestral population is shown. Each line represents an individual chromosome. B) The four scenarios considered are depicted with yellow stars represent mutations at the beneficial locus and red lines on the phylogenies when selection occurred. The scenarios are: 1) independent mutations, 2) independent selection on ancestral standing variation, 3) introgression, and 4) selection in the ancestral population. C) Panels for each of the four scenarios contain a cartoon representation of the haplotypic patterns surrounding the beneficial allele (black dot) in the two selected populations after fixation and a gene tree relating the populations at the beneficial allele. Pink regions in the haplotypes represent new mutations. D) Within- and between-population conacestry coefficients for the four scenarios are shown as a function of recombination distance from the beneficial allele. Both gene trees and coancestry coefficients were derived from simulations briefiy outlined in Appendix A.2.

Now, to build intuition, we briefly summarize the types of patterns we expect to observe under various sceanarios that result in an adaptive allele being fixed in two populations, as illustrated in Figure 2B. We focus on four cases: 1) independent selection on independent mutations for a beneficial allele at the same locus, 2) independent selection on an allele that was standing in the ancestor of the selected populations and independently in the daughter populations for a moderate amount of time, 3) introgression followed by independent selection, and 4) a complete sweep in the ancestral population of the two adapted populations prior to their splitting. The first three cases fall under our definition of convergent adaptation while the last would not due its lack of independence or replication. For each scenario, we focus on three signals: the haplotype patterns in our two adapted populations (Figure 2C), the gene tree at the beneficial allele for the two adapted populations plus an outgroup (Figure 2C), and the coancestry coefficients that our approach utilizes, which are another statement about the haplotypic similarity within and between populations, as a function of recombination distance from the locus experiencing selection (Figure 2D). The latter two signals shown were generated from simulations as outlined briefly in Appendix A.2.

The difference between the independent mutations scenario and the others is perhaps the most striking. Here, the beneficial allele arose on different haplotypes in the two adapted populations and, after fixation, we observe high levels of similarity within the two populations, but none between. This sharing within a population that experienced selection is due to the fact that variants linked the beneficial allele also increased in frequency. As we move further away from the selected locus, recombination is more likely to occur and this association breaks down. This is shown in the coancestry coefficients as a function of recombination distance from the selected site. In this case, there is an increase in coancestry within a selected population but no increase in coancestry (beyond neutral, shared population history) between the populations. We can also see this in the tree generated at the selected site. Here, all lineages within a population coalesce rapidly, while between them, the lineages do find a common ancestor faster than on the time-scale of typical, neutral lineages.

The other three scenarios have a key similarity. Now, there is some haplotypic similarity in both adapted populations since the beneficial allele has a single, shared origin. Still, there are features that differentiate these scenarios that may be useful to distinguish them. In the case that selection was on a variant that was standing in the ancestor of the adapted populations, there is a region of a shared, core haplotype between populations and, as a result, increased coancestry between the populations. However, we observe a faster decay in coancestry with increasing recombination distance between the populations than within a population. This is due to the fact that while the allele was standing independently in both populations prior to the onset of selection, recombination broke down the original haplotype on which the selected allele arose. Seeing a strong sweep within populations, but only a partially shared haplotype among the populations, is evidence that selection has independently dragged up the same variant from low frequency. In contrast, the final two scenarios have higher amounts of sharing both within and between the adapted populations. In the introgression case, we expect little to no differentiation in the haplotypes between the populations if migration occurs during the sweep. The gene tree depicts that all lineages with the beneficial allele coalesce rapidly at the selected site, regardless of which population they were sampled from. In the coancestry coefficients, there is now a slow decay in the within-population coancestries and a similar decay between populations. The final scenario, which does not represent convergent adaptation but rather a single, ancestral sweep, looks very similar to introgression since there is a lot of sharing in haplotypes within and between populations. The key difference is, if selection occurred sufficiently far in the past, we will observe new mutations independently arising in the adapted populations (indicated by the pink bars on the haplotypes). Specifically, in the tree we still observe all lineages at the selected site coalesce at the same time in the past, but now there are branches leading to our present-day lineages on which new mutations have occurred. Thus, to demonstrate that the beneficial allele swept in multiple populations, the between-population coalescent times should be much more recent than those in the rest of the genome. Observing very recent coalescence (i.e. short branch lengths) between and among populations in the swept region would allow us to rule out that the sweep is shared ancestrally or spread by neutral migration pressure. Such sweep patterns, that are independent among multiple populations, can provide evidence of adaptive convergence and the independent gain of information due to selection.

## Discussion

We emphasize the importance in carefully defining the questions we are trying to address in studying convergent adaptation. Convergent evolution has a wide range of overlapping definitions; it is obviously fine if our discussion does not line up with your definition (indeed one of the authors has varied in his definitions, Ralph and Coop, 2010, 2015b). However, we assert that when there has been convergent substitutions at a locus among taxa, it is of interest that selection favored their increase in frequency multiple times, no matter their origin.

There is a growing body of theory on when we should expect various modes of convergent adaptation among distantly-related populations (due to independent mutations Orr, 2005; Chevin et al., 2010) and among closely-related populations (due to independent mutations, standing variation, and the spread of alleles by migration Gordo and Campos, 2006; Ralph and Coop, 2010, 2015a,b; MacPherson and Nuismer, 2017; Paulose et al., 2018). Dissecting whether the same or different mutations gave rise to the adaptive alleles will help us gain insights into questions related to mutational target size or epistatic constraints, but these questions can be somewhat orthogonal to more basic questions about adaptation. As we learn more about repeated adaptation across populations, the need to differentiate convergent mutations from convergent allele frequency change will increase. For example, human populations have repeatedly adapted to different environments through changes in skin pigmentation. This convergent adaptation on the level of the phenotype comes from a mixture of selection on old standing variation, both derived and ancestral variants, and recent mutations (Norton et al., 2007; Edwards et al., 2010; Crawford et al., 2017). In the case of repeated evolution likely involving highly polygenic traits (such as human height and the convergent evolution of small body size; Perry and Dominy, 2009; Tucci et al., 2018), populations may be convergently adapting via very similar shared sets of alleles. This may reduce the independent information we can hope to glean about the genetics of height, but establishing that the phenotypic change occurred convergently is still important.

We demonstrated a potential way to quantify the impressiveness of various instances of convergent adaptation. One fair question is whether expressing these ideas in terms of information gained by selection and surprise add something new to the discussion. We ourselves are still a little unsure. After all, log-likelihood ratios are regularly computed as a means of summarizing support for various selection models over the null. However, usually the log-likelihood is viewed as a test statistic that should be compared to some null distribution, to see if the null model can be rejected. Indeed, we often tell ourselves that we should not be invested in the value of log-likelihood ratio as it is influenced by our sample size, etc. However, in calculating log-likelihoods of a particular allele being fixed in different populations, the likelihood ratio is a statement about the outcome of an evolutionary process and its replications across populations. Therefore, thinking about the log-likelihood of outcomes across populations as representations of the information gained by selection, at least under some models, focuses us on the view that natural selection is a force that drives the origination of the improbable, and in the case of convergent adaptation, its repeated origination. More work is needed to demonstrate that this is indeed useful and how it could be practically applied, but the idea itself may have merit.

In this article, we focus on how population-level genomic data, specifically from closely-related taxa, forces us to recognize new challenges in identifying convergent adaptation while also enables us to address these issues in new ways. Additionally, the sharing of characters among closely-related taxa brings its own set of opportunities. Repeated adaptation among recently diverged populations potentially allows us to use population genomics, genetic crosses, and association studies to map genotype to phenotype, as well as the option to perform experimental manipulations in a replicated populations across relatively uniform genetic backgrounds. Population genetic tools can be useful in helping to assess the two criteria necessary to have convergent adaptation. There are many existing approaches, that often nicely parallel those from phylogenetics, to test that changes we observe are unusual, relative to some null model. More work is needed to fully flesh out the potential of these methods, but these tests can be incorporated into existing frameworks allowing the field to move forward with more explicit tests for convergent adaptation. We are excited for what new data and tools can help us learn about convergence. Still, it is important, to emphasize that cases that meet these two criteria may help in identifying loci and traits that may be interesting to investigate further, but they might not be convergent adaptations on the phenotypic level. For example, two loci may have swept the same haplotype to fixation, we cannot be certain that the haplotypes, while sharing some alleles, do not also harbor different alleles conferring different phenotypic effects on fitness. Therefore, the hard work of linking the genotype and phenotype to fitness will remain an obstacle in describing convergent adaptation. However, the growing potential for identifying new cases of convergence in closely-related populations offers a rich seam for these efforts.

## Acknowledgements

We wish to thank Nick Barton, Jeremy Berg, Doc Edge, Emily Josephs, Matt Osmond, and members of the Coop lab for helpful discussions and feedback. We would also like to apologize to authors whose work we could not cite on this broad topic due to limited space. This work was supported by the National Science Foundation Graduate Research Fellowship awarded to KML (1148897) and by grants from the National Science Foundation (1353380) and the National Institute of General Medical Sciences of the National Institutes of Health (NIH R01 GM108779) awarded to GC.

## A Appendix

## A.1 Information

Kimura (1961) argued that Haldane‘s (1957) substitution load for a single selected allele is also related to the accumulation of genetic information. In this appendix, we discuss ideas about convergent allele frequency change in terms of information theory.

The information gain of the probability distribution *p*( ) over *q*( ) is defined as

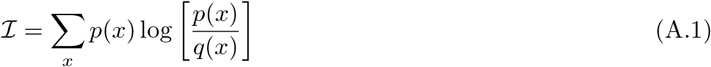

This is the log-likelihood of *x* averaged over the probability distribution *p*(). It measures how much more concentrated *p*( ) is on particular states in *x* compared to *q*(). It is also called the Kullback-Leibler divergence, or relative entropy of the first model to the second model.

We can write the gain of information under our selection model compared to a neutral model as

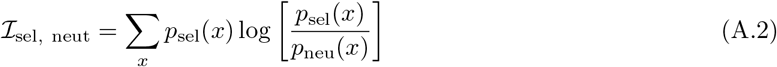

i.e. the log-likelihood ratio under our selected compared our neutral model averaged over outcomes of our selected model.

To see Kimura‘s (1961) argument, imagine an allele starting at frequency *x*_1_ in the population. If it were neutral, it would fix in the population with probability *x*_1_. If it is selected, it will fix with probability 1 (assuming *x*_1_≫ 1/(2*N_e_s*, where *N_e_* is the effective population size). Then, the gain of information under the selection model compared to the neutral model is

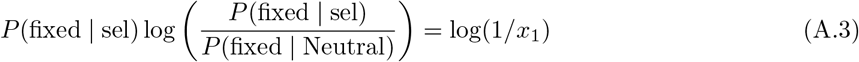

as originally outlined by Kimura (1961).

We can consider the gain of information in moving from frequency *x*_1_ to *x_2A_*, as in our population split case (Figure 1B). Under genetic drift, the probability that a neutral allele starting from frequency *x_1A_* ≤ *x_2A_* reaches frequency *x_2A_* before loss is *x_1_/x_2_*. An advantageous, additive allele achieves this change deterministically (again assuming (*x_1A_≫1/2_N_e_s_*). Therefore, we can write the gain of information of moving from frequency *x_1A_* to *x_2A_* under our selection model to our neutral model as

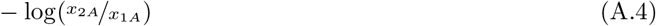

This shows that our selective deaths in achieving this frequency change (Equation 3) in the ancestral population may also be thought of as a selective gain in information. (Note in expressing things this way there is an ambiguity in that we state that the frequency at the moment the populations split is *x_2A_*. Therefore, a alternative formulation of the neutral probability may be the transitory stationary distribution at *x_2A_* of the neutral diffusion starting from frequency *x_1A_*. A different approach that would give us Equation A.4 is to think about the difference in the gain of information due to selection in the system when going from frequency *x*_1_ → 1 compared to going from *x*_2_ → 1.)

Thus, we can decompose the total information gain under the selection model of going from frequency *x_1A_* to *x_2A_* and then from *x_2A_* to fixation into *L_D_* + *L_A_*. Note that in doing this we have not specified the time to took selection to achieve this gain. If we have knowledge of the timing, this could be incorporated by using the allele-frequency transition probabilities for the neutral and selection model in Equation A.2. (Note that incorporating the timing would lead to the information gain to depend on the strength of selection.) Using transition probabilities would also allow us to incorporate genetic drift into the selection model, which would allow the amount of work done by selection to be averaged over likely values of the allele frequency change due to the combined actions of genetic drift and selection. Such considerations would need more work if this were to be useful in a practical setting.

Similarly, the total information accumulated in fixing the allele in populations 1 and 2 in our admixture example (Figure 1C) is

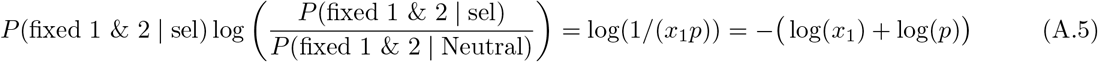

Therefore, one measure of adaptive convergence may be the additional information gained by selection independently in different populations. In the context of information theory, Equations 5 and 6 are called the coefficient of constraint which is a measure of how well we can predict the information in population 2 from that given to us by population 1.

**Surprise!** We can also formulate our results in terms of gain of information as the ‘surprise’ at our allelic state across populations. In information theory, the negative log probability of an outcome under a particular model is called the surprise (or the self information). The lower the probability of a particular outcome under a model, the greater the surprise when we see that outcome. Another nice feature, following from that of log probabilities, is that surprise is additive so you are doubly surprised if you see the same rare event twice independently.

As everything is done under a effective deterministic selection model (i.e. starting from a frequency ≫ 1/(2*N_e_s*), all of our statements about information gain only include the log probabilities under the neutral model. Therefore, they are the surprise at the outcome under the neutral model. Our measure of the work done by neutral probabilities can be decomposed into a product of conditional neutral, probabilities, e.g. Equation (A.5) and so a sum of log probabilities. This has the nice feature that we can ask how surprised we are at the convergence in a second population given a change in the first population.

## A.2 Coancestry coefficients

Knowledge of variation linked to a selected site may enable us to address questions of convergence that would not be possible if we only observed the state of the selected populations. If we observe sister populations sharing a selective event, we can ask if the selection was independent or if the selective event occurred in their ancestor. If the selected event is shared and the sweep is recent, we might expect the regions around the selected site to be more similar than if the sweeps happened independently and are truly convergent. We can formalize this in terms of coancestry between populations.

We define coancestry coefficient *f_ij_* as the probability that a pair of alleles sampled from populations *i* and *i* coalesce before the ancestral population of all sampled populations (see Lee and Coop, 2017, for more details). Thinking about the populations in Figure 1B, we can define the coancestry due to both neutral processes and selection for populations 1 and 2 as 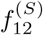 where ƒ_12_ alone specifies the coancestry between the populations due to drift and admixture. If the selective events occurred independently from new mutation of the beneficial allele in both populations, the coancestry between them at loci near the selected allele is simply what we expect under neutrality, *f*_12_.

The coancestry will increase if selection is on the same ancestral standing variant present in the ancestor of the populations, and is a function of the frequency of the standing variant, *g* (i.e. *x_2A_* in Figure 1B) and the amount of time, *t*, once populations 1 and 2 have split from their ancestor before selection started. Assuming the standing variant was previously neutral or was maintained in the population at some low frequency by balancing selection,

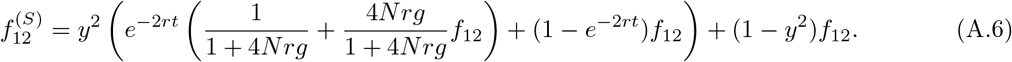

where *y^2^* represents the probability of both linked lineages failing to recombine off the beneficial allele. This can be approximated as e^-rt_s_^ where *t_s_* is the duration of the sweep phase. This increase in coancestry, that decays with distance from the selected site, is due to the fact the region around the beneficial allele looks more similar if the variant is shared. However, as the amount of time the beneficial allele is standing independently in the sister populations before selection occurs increases, we expect this similarity to decrease as recombination is occurring independently in the populations. See Lee and Coop (2017) for full derivation.

If the selective event is shared completely (i.e. the sweep occurs in the ancestor of populations 1 and 2 before they split), there is a further increase in coancestry. In this case the fraction of shared deaths is 1.

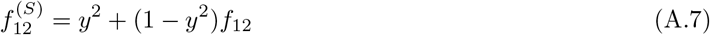

Even if the selective event is partially shared such that selection occurs in the ancestral population (for time *t*_1_) and continues independently in the daughter populations (for time *t*_2_), this takes the same form as (A.8) if *t*_1_ +*t*_2_ = *t*_s_.

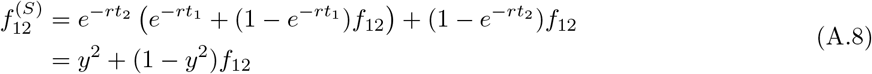

Therefore it may not be possible to distinguish cases where fraction of shared death is greater than zero such that selective event is shared completely or partially. However, it may be possible to detect if truly convergent i.e. there is no overlap in selected deaths.

**Simulation details.** We performed coalescent simulated under the four scenarios outline above using mssel, a modified version of ms (Hudson, 2002) that allows for the incorporation of selection at single site and stochastically generated allele frequency trajectories. See Lee and Coop (2017) for more details. These simulations were used to generate both coancestry coefficients averaged over 100 simulations and the gene trees. For all, we specified a 5% reduction in fitness for heterozygotes not carrying the beneficial allele and assumed a model of additivity. For the ancestral sweep model, we simulated a sweep that finished 0.1 coalescent generations in the past. For the standing variant model, we specified that the variant was standing at 1% frequency for 0.075 coalescent generations after the populations split. Additionally, for the migration model, we allowed for a fraction of 1 × 10^-4^ migrants per generation over the duration of the sweep from the source population into the recipient population.

## References

Adrion, J. R., M. W. Hahn, and B. S. Cooper (2015). Revisiting classic clines in drosophila melanogaster in the age of genomics. Trends in Genetics 31 (8), 434–444.

Arendt J. and D. Reznick (2008). Convergence and parallelism reconsidered: what have we learned about the genetics of adaptation? Trends in Ecology & Evolution 23(1), 26–32.

Avise J. C. and T. J. Robinson (2008). Hemiplasy: a new term in the lexicon of phylogenetics. Systematic Biology 57(3), 503–507.

Barton N. (2017). How does epistasis influence the response to selection? Heredity 118(1), 96–109.

Baum D. A. and A. Larson (1991). Adaptation reviewed: a phylogenetic methodology for studying character macroevolution. Systematic Biology 40(1), 1–18.

Berg, J., X. Zhang, and G. Coop (2017). Polygenic adaptation has impacted multiple anthropometric traits. bioRxiv.

Berg J. J. and G. Coop (2014, August). A population genetic signal of polygenic adaptation. PLoS Genetics 10 (8).

Berg J. J. and G. Coop (2015). A coalescent model for a sweep of a unique standing variant. Genetics 201 (2), 707–725.

Bierne N. (2010). The distinctive footprints of local hitchhiking in a varied environment and global hitchhiking in a subdivided population. Evolution 64 (11), 3254–3272.

Bonhomme, M., C. Chevalet, B. Servin, S. Boitard, J. M. Abdallah, S. Blott, and M. San Cristobal (2010). Detecting selection in population trees: the Lewontin and Krakauer test extended. Genetics.

Castoe, T. A., A. J. de Koning, H.-M. Kim, W. Gu, B. P. Noonan, G. Naylor, Z. J. Jiang, C. L. Parkinson, and D. D. Pollock (2009). Evidence for an ancient adaptive episode of convergent molecular evolution. Proceedings of the National Academy of Sciences 106(22), 8986–8991.

Chan, Y. F., M. E. Marks, F. C. Jones, G. Villarreal, M. D. Shapiro, S. D. Brady, A. M. Southwick, D. M. Absher, J. Grimwood, J. Schmutz, et al. (2010). Adaptive evolution of pelvic reduction in sticklebacks by recurrent deletion of a Pitx1 enhancer. Science 327(5963), 302–305.

Chevin, L.-M., G. Martin, and T. Lenormand (2010). Fisher’s model and the genomics of adaptation: restricted pleiotropy, heterogenous mutation, and parallel evolution. Evolution: International Journal of Organic Evolution 64 (11), 3213–3231.

Christin, P.-A., D. M. Weinreich, and G. Besnard (2010). Causes and evolutionary significance of genetic convergence. Trends in Genetics 26(9), 400–405.

Colosimo, P. F., K. E. Hosemann, S. Balabhadra, G. Villarreal, M. Dickson, J. Grimwood, J. Schmutz, R. M. Myers, D. Schluter, and D. M. Kingsley (2005). Widespread parallel evolution in sticklebacks by repeated fixation of ectodysplasin alleles. Science 307(5717), 1928–1933.

Coop, G., D. Witonsky, A. Di Rienzo, and J. K. Pritchard (2010). Using environmental correlations to identify loci underlying local adaptation. Genetics.

Cover T. M. and J. A. Thomas (2012). Elements of Information Theory. John Wiley & Sons.

Crawford, N. G., D. E. Kelly, M. E. B. Hansen, M. H. Beltrame, S. Fan, S. L. Bowman, E. Jewett, A. Ranciaro, S. Thompson, Y. Lo, S. P. Pfeifer, J. D. Jensen, M. C. Campbell, W. Beggs, F. Hormozdiari, S. W. Mpoloka, G. G. Mokone, T. Nyambo, D. Wolde Meskel, G. Belay, J. Haut, H. Rothschild, L. Zon, Y. Zhou, M. A. Kovacs, M. Xu, T. Zhang, K. Bishop, J. Sinclair, C. Rivas, E. Elliot, J. Choi, S. A. Li, B. Hicks, S. Burgess, C. Abnet, D. E. Watkins-Chow, E. Oceana, Y. S. Song, E. Eskin, K. M. Brown, M. S. Marks, S. K. Loftus, W. J. Pavan, M. Yeager, S. Chanock, and S. Tishkoff (2017). Loci associated with skin pigmentation identified in african populations. Science.

Duforet-Frebourg, N., K. Luu, G. Laval, E. Bazin, and M. G. Blum (2015). Detecting genomic signatures of natural selection with principal component analysis: application to the 1000 genomes data. Molecular Biology and Evolution 33(4), 1082–1093.

Edwards, M., A. Bigham, J. Tan, S. Li, A. Gozdzik, K. Ross, L. Jin, and E. J. Parra (2010, 03). Association of the oca2 polymorphism his615arg with melanin content in east asian populations: Further evidence of convergent evolution of skin pigmentation. PLOS Genetics 6(3), 1–8.

Felsenstein J. (1971). On the biological significance of the cost of gene substitution. The American Naturalist 105 (941), 1–11.

Felsenstein J. (1985). Phylogenies and the comparative method. The American Naturalist 125(1), 1–15.

Frank S. A. (2012). Natural selection. v. how to read the fundamental equations of evolutionary change in terms of information theory. Journal of Evolutionary Biology 25(12), 2377–2396.

Frichot, E., S. D. Schoville, G. Bouchard, and O. Francois (2013). Testing for associations between loci and environmental gradients using latent factor mixed models. Molecular Biology and Evolution 30(7), 1687–1699.

Fukushima, K., X. Fang, D. Alvarez-Ponce, H. Cai, L. Carretero-Paulet, C. Chen, T.-H. Chang, K. M. Farr, T. Fujita, Y. Hiwatashi, et al. (2017). Genome of the pitcher plant cephalotus reveals genetic changes associated with carnivory. Nature Ecology & Evolution 1 (3).

Futuyma D. (2013). Evolution. Sinauer.

Galinsky, K. J., G. Bhatia, P.-R. Loh, S. Georgiev, S. Mukherjee, N. J. Patterson, and A. L. Price (2016). Fast principal-component analysis reveals convergent evolution of ADH1B in Europe and East Asia. The American Journal of Human Genetics 98 (3), 456–472.

Givnish T. J. (2015). New evidence on the origin of carnivorous plants. Proceedings of the National Academy of Sciences 112(1), 10–11.

Gordo I. and P. R. Campos (2006). Adaptive evolution in a spatially structured asexual population. Genetica 127(1-3), 217.

Goudet, J., T. Kay, and B. S. Weir (2018). How to estimate kinship. Molecular Ecology.

Gould S. J. and E. S. Vrba (1982). Exaptationa missing term in the science of form. Paleobiology 8(1), 4–15.

Guerrero R. F. and M. W. Hahn (2018). Quantifying the risk of hemiplasy in phylogenetic inference. Proceedings of the National Academy of Sciences.

Günther T. and G. Coop (2013). Robust identification of local adaptation from allele frequencies. Genetics.

Hadfield J. and S. Nakagawa (2010). General quantitative genetic methods for comparative biology: phylogenies, taxonomies and multi-trait models for continuous and categorical characters. Journal of Evolutionary Biology 23(3), 494–508.

Hahn M. W. and L. Nakhleh (2016). Irrational exuberance for resolved species trees. Evolution 70(1), 7–17.

Haldane J. B. S. (1957, Dec). The cost of natural selection. Journal of Genetics 55(3), 511.

Hancock, A. M., D. B. Witonsky, A. S. Gordon, G. Eshel, J. K. Pritchard, G. Coop, and A. Di Rienzo (2008). Adaptations to climate in candidate genes for common metabolic disorders. PLoS genetics 4 (2).

Harmon L. (2018).Phylogenetic Comparative Methods learning from trees.

Harvey, P. H., M. D. Pagel, et al. (1991). The comparative method in evolutionary biology, Volume 239. Oxford University Press.

Housworth, E. A., E. P. Martins, and M. Lynch (2004). The phylogenetic mixed model. The American Naturalist 163(1), 84–96.

Hudson R. R. (2002). Generating samples under a Wright-Fisher neutral model of genetic variation. Bioinformatics 18, 337–338.

Iwasa Y. (1988). Free fitness that always increases in evolution. Journal of Theoretical Biology 135(3), 265–281.

Josephs, E. B., J. J. Berg, J. Ross-Ibarra, and G. Coop (2018). Detecting adaptive differentiation in structured populations with genomic data and common gardens. bioRxiv, 368506.

Kaplan, N. L., R. R. Hudson, and C. H. Langley (1989, December). The “hitchhiking effect” revisited. Genetics 123, 887–899.

Karhunen, M., J. Merila, T. Leinonen, J. Cano, and O. Ovaskainen (2013). DRIFTSEL: an R package for detecting signals of natural selection in quantitative traits. Molecular Ecology Resources 13(4), 746–754.

Kim Y. and T. Maruki (2011). Hitchhiking effect of a beneficial mutation spreading in a subdivided population. Genetics 189(1), 213–226.

Kimura M. (1961). Natural selection as the process of accumulating genetic information in adaptive evolution. Genetics Research 2(1), 127–140.

Lee K. M. and G. Coop (2017). Distinguishing among modes of convergent adaptation using population genomic data. Genetics 207(4), 1591–1619.

Leinonen, T., R. S. McCairns, R. B. O’hara, and J. Merila (2013). Qst-Fst comparisons: evolutionary and ecological insights from genomic heterogeneity. Nature Reviews Genetics 14 (3), 179.

Losos J. B. (2011). Convergence, adaptation, and constraint. Evolution 65(7), 1827–1840.

Lynch M. (1991). Methods for the analysis of comparative data in evolutionary biology. Evolution 45(5), 1065–1080.

Machado, H. E., A. O. Bergland, K. R. OBrien, E. L. Behrman, P. S. Schmidt, and D. A. Petrov (2016). Comparative population genomics of latitudinal variation in Drosophila símulans and Drosophila melanogaster. Molecular ecology 25(3), 723–740.

MacPherson A. and S. Nuismer (2017). The probability of parallel genetic evolution from standing genetic variation. Journal of Evolutionary Biology 30(2), 326–337.

Mahler, D. L., T. Ingram, L. J. Revell, and J. B. Losos (2013). Exceptional convergence on the macroevolutionary landscape in island lizard radiations. Science 341 (6143), 292–295.

Martin A. and V. Orgogozo (2013). The loci of repeated evolution: a catalog of genetic hotspots of phenotypic variation. Evolution 67(5), 1235–1250.

Maynard Smith, J. (1968). “Haldane’s dilemma” and the rate of evolution. Nature 219(5159), 1114–1116.

Maynard Smith, J. and J. Haigh (1974, February). The hitch-hiking effect of a favourable gene. Genet Res 23(1), 23–35.

Mendes, F. K., J. A. Fuentes-Gonzalez, J. G. Schraiber, and M. W. Hahn (2018). A multispecies coalescent model for quantitative traits. eLife 7.

Mustonen V. and M. Lassig (2010). Fitness flux and ubiquity of adaptive evolution. Proceedings of the National Academy of Sciences 107(9), 4248–4253.

Norton, H. L., R. A. Kittles, E. Parra, P. McKeigue, X. Mao, K. Cheng, V. A. Canfield, D. G. Bradley, B. McEvoy, and M. D. Shriver (2007). Genetic evidence for the convergent evolution of light skin in europeans and east asians. Molecular Biology and Evolution 24 (3), 710–722.

Orr H. A. (2005). The probability of parallel evolution. Evolution 59(1), 216–220.

Ovaskainen, O., M. Karhunen, C. Zheng, J. M. C. Arias, and J. Merila (2011). A new method to uncover signatures of divergent and stabilizing selection in quantitative traits. Genetics 189(2), 621–632.

Paulose, J., J. Hermisson, and O. Hallatschek (2018). Spatial soft sweeps: patterns of adaptation in populations with long-range dispersal. bioRxiv, 299453.

Peck J. R. and D. Waxman (2010). Is life impossible? Information, sex, and the origin of complex organisms. Evolution: Internationa! Journal of Organic Evolution 64 (11), 3300–3309.

Perry G. H. and N. J. Dominy (2009). Evolution of the human pygmy phenotype. Trends in Ecology & Evolution 24 (4), 218–225.

Peter, B. M., E. Huerta-Sanchez, and R. Nielsen (2012). Distinguishing between selective sweeps from standing variation and from a de novo mutation. PLoS Genetics 8 (10).

Pickrell J. K. and J. K. Pritchard (2012). Inference of population splits and mixtures from genome-wide allele frequency data. PLoS Genetics 8(11).

Racimo F. (2016). Testing for ancient selection using cross-population allele frequency differentiation. Genetics 202(2), 733–750.

Racimo, F., J. J. Berg, and J. K. Pickrell (2018). Detecting polygenic adaptation in admixture graphs. Genetics.

Ralph P. L. and G. Coop (2010). Parallel adaptation: one or many waves of advance of an advantageous allele? Genetics.

Ralph P. L. and G. Coop (2015a). Convergent evolution during local adaptation to patchy landscapes. PLoS Genetics 11 (11).

Ralph P. L. and G. Coop (2015b). The role of standing variation in geographic convergent adaptation. The America/n Naturalist 186(S1), S5–S23.

Refoyo-Martínez, A., R. R. da Fonseca, K. Halldórsdottir, E. iÁrnason, T. Mailund, and F. Racimo (2018). Identifying loci under positive selection in complex population histories. bioRxiv.

Roesti, M., S. Gavrilets, A. P. Hendry, W. Salzburger, and D. Berner (2014). The genomic signature of parallel adaptation from shared genetic variation. Molecular Ecology 23(16), 3944–3956.

Rosenblum, E. B., C. E. Parent, and E. E. Brandt (2014). The molecular basis of phenotypic convergence. Annual Review of Ecology, Evolution, and Systematics 45, 203–226.

Santiago E. and A. Caballero (2005). Variation after a selective sweep in a subdivided population. Genetics 169, 475–483.

Sedghifar, A., P. Saelao, and D. J. Begun (2016). Genomic patterns of geographic differentiation in Drosophila simulans. Genetics 202(3), 1229–1240.

Shannon C. E. (1948). A mathematical theory of communication. Bell System Technical Journal. 27(3), 3–55.

Shapiro, M. D., M. E. Marks, C. L. Peichel, B. K. Blackman, K. S. Nereng, B. Joónsson, D. Schluter, and D. M. Kingsley (2004). Genetic and developmental basis of evolutionary pelvic reduction in threespine sticklebacks. Nature 428(6984), 717.

Slatkin M. and T. Wiehe (1998). Genetic hitch-hiking in a subdivided population. Genetical Research 71 (02), 155–160.

Stayton C. T. (2008). Is convergence surprising? An examination of the frequency of convergence in simulated datasets. Journal of Theoretical Biology 252(1), 1–14.

Stern D. L. (2013). The genetic causes of convergent evolution. Nature Reviews Genetics 14 (11), 751.

Stone, G. N., S. Nee, and J. Felsenstein (2011). Controlling for non-independence in comparative analysis of patterns across populations within species. Philosophical Transactions of the Royal Society of London B: Biological Sciences 366(1569), 1410–1424.

Storz J. F. (2016). Causes of molecular convergence and parallelism in protein evolution. Nature Reviews Genetics 17(4), 239.

Sved J. (1968). Possible rates of gene substitution in evolution. The American Naturalist 102(925), 283–293.

Thomas G. W. and M. W. Hahn (2015). Determining the null model for detecting adaptive convergence from genomic data: a case study using echolocating mammals. Molecular Biology and Evolution 32(5), 1232–1236.

Thorogood, C. J., U. Bauer, and S. J. Hiscock (2018). Convergent and divergent evolution in carnivorous pitcher plant traps. New Phytologist 217(3), 1035–1041

Tucci, S., S. H. Vohr, R. C. McCoy, B. Vernot, M. R. Robinson, C. Barbieri, B. J. Nelson, W. Fu, G. A. Purnomo, H. Sudoyo, et al. (2018). Evolutionary history and adaptation of a human pygmy population of flores island, indonesia. Science 361 (6401), 511–516.

Wain-wright P. C. (2007). Functional versus morphological diversity in macroevolution. Annual Review of Ecology, Evolution, and Systematics 38(1), 381–401.

Wake, D. B., M. H. Wake, and C. D. Specht (2011). Homoplasy: from detecting pattern to determining process and mechanism of evolution. Science 331 (6020), 1032–1035.

Watkins C. (2002). The channel capacity of evolution: ultimate limits on the amount of information maintainable in the genome. In Proceedings of the Third Internationa! Conference on Bioinformatics of Genome Regulation and Structure, Volume 2, pp. 58–60.

Welch J. J. and C. D. Jiggins (2014). Standing and flowing: the complex origins of adaptive variation. Molecular ecology 23(16), 3935–3937.

Whitlock M. C. (2008). Evolutionary inference from Qst. Molecular Ecology 17(8), 1885–1896.

Yeaman, S., A. C. Gerstein, K. A. Hodgins, and M. C. Whitlock (2018). Quantifying how constraints limit the diversity of viable routes to adaptation. PLOS Genetics 14 (10), 1–25.

Zhang J. and S. Kumar (1997). Detection of convergent and parallel evolution at the amino acid sequence level. Molecular Biology and Evolution 14 (5), 527–536.

